# A variant of the student’s *t*-test for data of varying reliability

**DOI:** 10.1101/525774

**Authors:** Suril B. Sheth, Bhavin R. Sheth

## Abstract

The student’s *t*-test has been a workhorse of statistical testing and is used to determine if two sets of sampled data are significantly different from one another, in a statistical sense. The samples of the data may be individual samples or the means – or some overall summary statistic – of independently acquired subsets of data (e.g. data from individual observers, neurons, or baseball games). The various subsets of data acquired that go into computing the t-statistic are likely to be of differing reliability on account of either different variances or of different numbers of subsamples corresponding to each subset; while all data are given equal weight in a standard *t*-test, the variation in data reliability across subsets of data needs to be accounted for. Solutions based on mixed model methods and Monte Carlo simulations exist, which do factor data reliability in computing statistics. However, no such extension exists for the ubiquitous student’s *t*-test. Our proposal is a novel variant of the students *t*-test that incorporates these issues and adopts a simple but effective alteration in the design that accounts for differing levels of data reliability. Specifically, we weighted each data subset by the inverse of the variance of the data contained therein, a measure that has been used in studies of Bayesian cue combination, or, in the absence of information about variance, by the relative proportion of the overall data contained in the subset. The changes proposed here extend the applicability of the student’s *t*-test to a wider array of data sets.

## Introduction

Data acquisition is a process in which the best-planned experiments can often get waylaid by random events, which affect the underlying reliability of the data. One way around this issue is to simply discard data that are not of the highest reliability and work with limited data that we know are of extremely high reliability. However, one might argue that data acquisition is a process that is inherently noise-ridden and unreliable, working with data that are not perfectly reliable is a fact of life, discarding data that are at least partly reliable reduces the effect size and statistical power, and data that are at least partly reliable also provide some useful information and must not be discarded.

Consider the following example from signal processing. A voltmeter measures the voltage in an electrical circuit, and the requirement is for the voltage to be above a prescribed value. The signal is measured multiple (albeit limited) number of times to get a more accurate value of signal voltage; each time the analog signal is digitized by one of a bank of analog to digital converters (ADC); the resolution, i.e. number of bits N, of each ADC differs. The signal to noise ratio (SNR) of the quantized digital signal varies linearly as a function of N. The average digital voltage computed needs to account for the differences in resolution among the different ADCs before being statistically compared with the prescribed threshold.

Moreover, some experiments are natural, “real world” experiments that cannot be controlled and inevitably generate data that have different amounts of reliability. Here are two examples from education and neuroscience (one can think of analogous examples from other areas).

Suppose the state introduces a new standardized test and a school has recently instructed its teachers to teach to the test, whereas a second neighboring school within the same district has done no such thing. In order to find out how well the teachers have succeeded in teaching to the test, one needs to compare mean test scores of the students in the teachers’ classes in each of the two districts. Classes can be and are usually of different sizes – and this is not typically under the teacher’s control; the reliability of the mean test score obtained from a class size of 10 students versus a class size of 50 students has to be accounted for.

A second example is from neuroscience. Researchers want to examine the relationship between consolidation of overnight learning and power contained in the delta band of frequencies during slow-wave sleep (SWS). In order to find out delta power in SWS, one computes the power contained in every single 30 second long epoch of SWS and then take the average across all such epochs (alternatively, one can combine all the 30 second epochs of SWS across all sleep cycles and then compute the power). Here, different individuals have different amounts of SWS and the variation in SWS amounts can be large and is not under the experimenter’s control. This variation in SWS amounts across individuals is a factor that has to be accounted for in developing statistical tests. Thus, when acquired under different conditions with different levels of reliability or averaged over different numbers of repetitions, data are not equally reliable and the differing levels of reliability is an important factor that must be taken into consideration in conducting statistical tests.

Therefore, a reasonable alternative is to retain the partly reliable data and weigh the reliability of the data relative to the reliability of other data in statistical tests. There are several avenues available including mixed model statistics and Monte Carlo methods.

The student’s *t*-test is arguably the most commonly used test of statistical significance, and clearly the first one taught in AP statistics classes in the country. However, as it stands, the *t*-test cannot account for the real world differences in reliability. Here, we offer a modification to the *t*-test that takes into account the reliability of disparate data.

## Methods

The t-statistic is defined by t = Z /s where Z and s are functions of the data; Z is a measure of the difference in the means in units of standard deviation of the sample; s is the standard error of the mean. More generally, 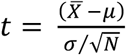 where 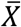 is the sample mean of a sample *X*_1_, *X*_2_,…. *X*_*N*_ of size N, *μ* is the population mean, and *σ* is the population standard deviation of the data. Samples *X*_1_, *X*_2_,…. *X*_*N*_ can be individual samples or may themselves be means of independent samples of the data, i.e. each *X*_*i*_ is the mean of samples 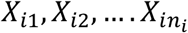, where *n*_*i*_ is the number of samples. The formula for the population standard deviation is as follows

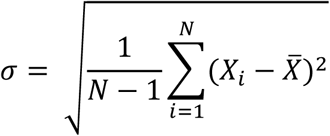

a)In the latter case, the reliabilities of the individual samples are likely to differ, which is otherwise ignored in the conventional formula for the t-statistic. We can define reliability *r*_*i*_ of the i^th^ sample as the inverse of the variance 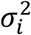 of the i^th^ sample data, i.e.

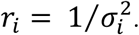

Note that [1-3] have used a similar Bayesian formula for weighting the relative reliabilities of sources. We define a new reweighted mean 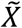, which is defined as the following:

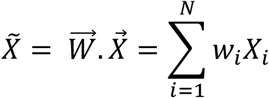

Where

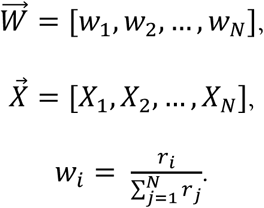

Note that

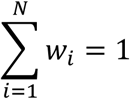

The revised weighted population standard deviation 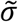 is as follows

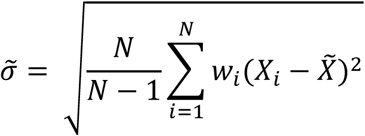

The newly revised t-statistic 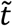 is now given by

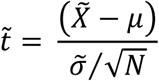

b) In cases where the individual data points 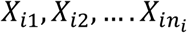 are not recorded or available for some reason, we designed a minor variant in which the reliability of each datum *X*_*i*_ is (proportional to) the number of the number of data points *n*_*i*_ that went into the calculation of *X*_*i*_, i.e.

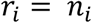

The remaining calculations for 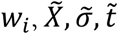, are identical to a).

## Results

Here, we analyze data inspired by research and demonstrate a numerical application of the *t*-test variant to the data. Liu and Sheth [4] studied the response of the brain to pure tones (1 kHz) of varying intensity (53/58/63 dB SPL) in wake and in stage II sleep. In particular, the study focused on the P200 component of the electroencephalography (EEG) response, and asked if the amplitude of the P200 differed between the two states for different sound intensities. Six subjects participated in the experiment. For each individual subject, the mean P200 amplitude was computed for each state by averaging data from 300 trials / state. The difference in mean P200 amplitudes in wake and SII sleep (P200_wake_ – P200_SII_) across all six subjects can be compared with zero difference in a classical paired *t*-test paradigm.

Using a classic paired *t*-test paradigm, for the 53 dB sound intensity, the calculations yielded the following results summarized in Table 2:

**Table 1:** P200 ERP amplitudes in response to different sound intensities

| Sound intensity | 53 dB |  |  | 58 dB |  |  | 63 dB |  |
| --- | --- | --- | --- | --- | --- | --- | --- | --- |
| Subject | Wake ($\mu\text{V}$) | SII ($\mu\text{V}$) | | Wake ($\mu\text{V}$) | SII ($\mu\text{V}$) | | Wake ($\mu\text{V}$) | SII ($\mu\text{V}$) |
| S1 | 6.67 | 2.81 |  | 6.57 | 7.38 |  | 11.87 | 7.38 |
| S2 | 6.61 | 3.55 |  | 9.64 | 2.94 |  | 7.42 | 11.95 |
| S3 | 5.06 | 2.85 |  | 5.20 | 5.15 |  | 11.17 | 5.82 |
| S4 | 6.22 | 3.01 |  | 7.16 | 4.83 |  | 10.94 | 8.24 |
| S5 | 4.09 | 5.22 |  | 6.88 | 6.29 |  | 6.36 | 4.67 |
| S6 | 6.44 | 4.82 |  | 8.75 | 6.75 |  | 5.96 | 9.88 |

**Table 2:** Calculations in classic paired *t*-test paradigm

| Sound intensity | 53 dB | 58 dB | 63 dB |
| --- | --- | --- | --- |
| | $\bar{X}_{\text{Wake-SII}} = 2.14 \mu\text{V}$ | $\bar{X}_{\text{Wake-SII}} = 1.81 \mu\text{V}$ | $\bar{X}_{\text{Wake-SII}} = 0.96 \mu\text{V}$ |
| | $\sigma_{\text{Wake-SII}} = 1.78 \mu\text{V}$ | $\sigma_{\text{Wake-SII}} = 2.67 \mu\text{V}$ | $\sigma_{\text{Wake-SII}} = 4.22 \mu\text{V}$ |
| | $t(5) = \frac{2.14}{1.78/\sqrt{6}} = 2.94$ | $t(5) = \frac{1.81}{2.67/\sqrt{6}} = 1.66$ | $t(5) = \frac{0.96}{4.22/\sqrt{6}} = 0.56$ |
| | $p = 0.032$ | $p = 0.158$ | $p = 0.602$ |

The above calculations did not take into account the variable reliabilities of the individual subject data. The numbers of trials during which the individual is awake and is in SII sleep are given below in Table 3. As can be seen, the numbers of trials vary across subjects.

The variant proposed here takes the variation in numbers of trials (or variance, for that matter, not shown here) into account in a new *t*-test paradigm. The results of the calculations are given in Table 4 below.

| Number of trials | 53 dB |  |  | 58 dB |  |  | 63 dB |  |
| --- | --- | --- | --- | --- | --- | --- | --- | --- |
| Subject | Wake | SII |  | Wake | SII |  | SII | Wake |
| S1 | 620 | 715 |  | 611 | 565 |  | 309 | 323 |
| S2 | 524 | 389 |  | 405 | 312 |  | 731 | 497 |
| S3 | 320 | 509 |  | 529 | 416 |  | 420 | 334 |
| S4 | 458 | 334 |  | 334 | 377 |  | 331 | 302 |
| S5 | 371 | 409 |  | 312 | 459 |  | 571 | 402 |
| S6 | 315 | 402 |  | 354 | 423 |  | 317 | 503 |

**Table 4:** Calculations in new paired *t*-test paradigm

| Sound intensity | 53 dB | 58 dB | 63 dB |
| --- | --- | --- | --- |
| | $\tilde{X}_{\text{Wake-SII}} = 2.35 \mu\text{V}$ | $\tilde{X}_{\text{Wake-SII}} = 1.49 \mu\text{V}$ | $\tilde{X}_{\text{Wake-SII}} = 0.29 \mu\text{V}$ |
| | $\tilde{\sigma}_{\text{Wake-SII}} = 1.76 \mu\text{V}$ | $\tilde{\sigma}_{\text{Wake-SII}} = 2.61 \mu\text{V}$ | $\tilde{\sigma}_{\text{Wake-SII}} = 4.34 \mu\text{V}$ |
| | $\tilde{t}(5) = \frac{2.35}{1.76/\sqrt{6}} = 3.26$ | $\tilde{t}(5) = \frac{1.49}{2.61/\sqrt{6}} = 1.39$ | $\tilde{t}(5) = \frac{0.29}{4.34/\sqrt{6}} = 0.16$ |
| | $p = 0.022$ | $p = 0.222$ | $p = 0.878$ |

Here, the re-weighting of data according to relative reliability changed the t-statistics and p-values slightly – t statistic changed from 2.94 to 3.26, 1.66 to 1.39, and 0.56 to 0.16 for53 dB, 58 dB, and 63 dB sound intensities respectively. The changes were small but more in line with the underlying data and the data acquisition process that produced them.

Our proposed variant can also be used in an unpaired setting as well. For this, we assume, for illustrative purposes, that the data for wake and SII were acquired from different subjects, which calls for an unpaired *t*-test for statistical testing. The t-statistic used for an independent two-samples test when both distributions have the same variance - here, the sample size is small enough that the equal variance assumption is not violated – is known to be given by the following formula 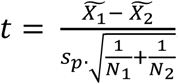 where *s_p_* is the pooled standard deviation of the two samples and, 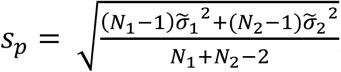 *N*_*1*_ is the size of the first sample (wake data, in the present case), and *N*_*2*_ is the size of the second sample (SII data, here)[5]. The calculations are given in Table 5 below.

**Table 5:** Calculations in new unpaired *t*-test

| 53 dB | 58 dB | 63 dB |
| --- | --- | --- |
| $\vec{W}_{Wake}$<br>$= [.24, .20, .12, .18, .14, .12]$<br>$\vec{W}_{SII}$<br>$= [.26, .14, .18, .12, .15, .15]$ | $\vec{W}_{Wake}$<br>$= [.24, .16, .21, .13, .12, .13]$<br>$\vec{W}_{SII}$<br>$= [.22, .12, .16, .15, .18, .17]$ | $\vec{W}_{Wake}$<br>$= [.12, .27, .16, .12, .21, .12]$<br>$\vec{W}_{SII}$<br>$= [.14, .21, .14, .13, .17, .21]$ |
| $\tilde{X}_{Wake} = 5.99 \mu V$<br>$\tilde{X}_{SII} = 3.60 \mu V$ | $\tilde{X}_{Wake} = 7.19 \mu V$<br>$\tilde{X}_{SII} = 5.80 \mu V$ | $\tilde{X}_{Wake} = 8.56 \mu V$<br>$\tilde{X}_{SII} = 8.30 \mu V$ |
| $\tilde{\sigma}_{Wake} = 1.00 \mu V$<br>$\tilde{\sigma}_{SII} = 1.05 \mu V$<br>$s_p = \sqrt{\frac{((6-1)*1.00^2 + (6-1)*1.05^2)}{6+6-2}}$<br>$= 1.03$ | $\tilde{\sigma}_{Wake} = 1.63 \mu V$<br>$\tilde{\sigma}_{SII} = 1.53 \mu V$<br>$s_p = \sqrt{\frac{((6-1)*1.63^2 + (6-1)*1.53^2)}{6+6-2}}$<br>$= 1.58$ | $\tilde{\sigma}_{Wake} = 2.50 \mu V$<br>$\tilde{\sigma}_{SII} = 2.81 \mu V$<br>$s_p = \sqrt{\frac{((6-1)*2.50^2 + (6-1)*2.81^2)}{6+6-2}}$<br>$= 2.66$ |
| $\tilde{t}(5) = \frac{5.99-3.60}{1.03 * \sqrt{\frac{1}{6} + \frac{1}{6}}} = 4.04$ | $\tilde{t}(5) = \frac{7.19-5.80}{1.58 * \sqrt{\frac{1}{6} + \frac{1}{6}}} = 1.53$ | $\tilde{t}(5) = \frac{8.56-8.30}{2.66 * \sqrt{\frac{1}{6} + \frac{1}{6}}} = 0.16$ |
| $p = 0.010$ | $p = 0.187$ | $p = 0.876$ |

## Discussion

The student’s *t*-test has had a rich history. Since being designed by William Sealy Gossett for determining if two sets of data are significantly different from each other [6], it has been extended to other important cases such as unpaired (independent) and paired (correlated) two-sample *t*-tests, tests for when the size of the two samples is not equal, Welch’s *t*-tests for when the two samples have unequal variance and/or unequal sample size [7], and a multivariate Hotelling’s test for multiple, often correlated, measures within the same sample [8].

Here, we propose a heteroaxiopistíc (hetero = different, axiopistía = reliability) variant of the student’s *t*-test so as to factor the relative reliabilities of the samples in the computation of the t-statistic. Reliability of data is an important variable and regularly factors into statistical testing and with the introduction of said variant here, data reliability can now factor into the popular students *t*-test as well. The methods described above take into account reliability of one set of samples, on a one-sample *t*-test. In the event of a comparison between two sets of samples, such as in a two sample *t*-test, the exact same methods as detailed above can be replicated for a second set of samples.

There are alternative powerful solutions to the problem of combining fixed and random effects in a mixed model [9]. Hierarchical linear effects models assume that data that are being analyzed are drawn from a hierarchy of different populations whose differences relate to that hierarchy [10]. Fixed and random effects typically refer to the population average and subject-specific effects, respectively and these effects are modeled using a classical matrix notation and fitted using an expectation maximization algorithm [11] where the variance components are treated as nuisance parameters that one does not care about but has to nonetheless account for. There are clear benefits to a mixed model approach, including the fact that it can deal with missed values or measurements in the data with remarkable ease - a likely possibility in data acquisition, and mixed models have justifiably formed the basis for a large amount of statistical research in recent years. The present approach to designing a heteroaxiopistíc variant of the student’s *t*-test provides, under certain conditions, a simpler and yet effective alternative that is more accessible to the end-user than more sophisticated mixed model approaches.

